# Effects of hyperinsulinemia on pancreatic cancer development and the immune microenvironment revealed through single-cell transcriptomics

**DOI:** 10.1101/2021.03.10.434504

**Authors:** Anni M.Y. Zhang, Twan J.J. de Winter, Su Wang, Stephane Flibotte, Yiwei Bernie Zhao, Xiaoke Hu, Hong Li, David F. Schaeffer, James D. Johnson, Janel L. Kopp

## Abstract

Hyperinsulinemia is independently associated with increased risk and mortality of pancreatic cancer. We recently reported that a ∼50% reduction in pancreatic intraepithelial neoplasia (PanIN) pre-cancerous lesions in mice could be achieved with reduced insulin production. However, only female mice remained normoglycemic and only the gene dosage of rodent-specific *Ins1* alleles was tested in our previous model. Moreover, we did not delve into the molecular and cellular mechanisms associated with modulating hyperinsulinemia. Here, we studied PanIN lesion development in both male and female *Ptf1a*^CreER^;*Kras*^LSL-G12D^ mice lacking the rodent specific *Ins1* gene, and possessing one or two alleles of the wild-type *Ins2* gene to modulate insulin production. High-fat diet induced hyperinsulinemia was transiently and modestly reduced, without affecting glucose tolerance, in male and female mice with only one allele of *Ins2*. Genetic reduction of insulin production resulted in mice with a tendency for less PanIN and acinar-to-ductal metaplasia (ADM) lesions. Using single-cell transcriptomics, we found hyperinsulinemia affected multiple cell types in the pancreas, with the most statistically significant effects on local immune cell populations, which were highly represented in our analysis. Specifically, hyperinsulinemia modulated pathways associated with protein translation, MAPK-ERK signaling, and PI3K-AKT signaling, which were changed in epithelial cells and subsets of immune cells. These data suggest a role for the immune microenvironment in hyperinsulinemia-driven PanIN development. Together with our previous work, we propose that mild suppression of insulin levels may be useful in preventing pancreatic cancer by acting on multiple cell types.

## Introduction

Pancreatic cancer, specifically pancreatic ductal adenocarcinoma (PDAC), is projected to become the 2^nd^ leading cause of cancer death in the next decade. Representing an estimated 2.5% of all new cancer cases, PDAC has a poor 5-year survival rate (1). Smoking, pancreatitis, family history, obesity and type 2 diabetes are risk factors for PDAC (2-6). Scientists predict obesity to overtake smoking and become the leading preventable cause of cancer (7), so efforts to understand the roles of diet and lifestyle in cancer risk and prevention strategies are expanding, as are efforts to determine the underlying pathophysiological mechanisms that mediate obesity-driven risk. Obesity and type 2 diabetes are usually accompanied by metabolic disorders like hyperinsulinemia, hyperglycemia, increased inflammation, and dyslipidemia, each of which are candidate factors that may contribute to the associated increase of cancer morbidity and mortality (8-12).

Hyperinsulinemia can be defined as excess circulating insulin, which is more than what is required for maintaining glucose homeostasis (12). Excess insulin is associated with an increased risk of cancer death that can be independent of obesity (13). Hyperinsulinemia has also been shown to be associated with increased incidence of PDAC and increased cancer mortality rate (14-18). Complementing these epidemiological studies, our recent *in vivo* animal study showed that hyperinsulinemia played a causal role in PDAC initiation in the context of a known hyperinsulinemia-inducing high fat diet (HFD) (19). Specifically, we found that *Ptf1a*^CreER^;*Kras*^LSL-G12D^ female mice, a commonly used PDAC mouse model, with ∼50% reduction in fasting insulin (*Ins1*^+/-^;*Ins2*^-/-^ compared to *Ins1*^+/+^;*Ins2*^-/-^), but no difference in glucose homeostasis, had a ∼50% reduction in PanIN lesions and fibrogenesis. Unfortunately, the experimental male mice were unable to tolerate having only a single allele of *Ins1* and rapidly became hyperglycemic. Nevertheless, these studies were the first to directly demonstrate that endogenous hyperinsulinemia contributes to development of any cancer (19).

The parental imprinting, gene structure and tissue distribution of murine *Ins2* gene is similar to human *INS* gene (20-23). It is therefore important to determine if PanIN initiation would also be affected by reducing *Ins2* gene dosage in *Ptf1a*^CreER^;*Kras*^LSL-G12D^ mice. Moreover, *Ins1* contributes to only ∼1/3 of secreted insulin; this means that *Ins1*^+/-^;*Ins2*^-/-^ mice have the lowest amount of insulin compatible with survival, with only female mice remaining consistently normoglycemic in modern mouse housing facilities. In the current study, we addressed the effect of hyperinsulinemia on PDAC development in both sexes by generating *Ptf1a*^CreER^;*Kras*^LSL-G12D^ mice with the *Ins1*^-/-^;*Ins2*^+/+^ or *Ins1*^-/-^;*Ins2*^+/-^ genotypes. Comparing *Ins1*^-/-^;*Ins2*^+/-^ experimental mice to *Ins1*^-/-^;*Ins2*^+/+^ controls allowed us to compare males and females side-by-side. The current study also provided an opportunity to examine pancreatic single-cell transcriptomics and gain insights into the molecular mechanisms involved in the complex cellular landscape of PDAC initiation under hyperinsulinemic conditions.

## Experimental procedures

### Mice

University of British Columbia Animal Care Committee in accordance with Canadian Council for Animal Care guidelines approved all animal experiments. Mice were kept at the University of British Columbia Modified Barrier Facility (MBF) as previously described in details (19). *Ptf1a*^CreER/WT^, *Kras*^LSL-G12D/WT^, *Ins1*^-/-^, and *Ins2*^-/-^ mice have been previously described (19, 22, 24-27). *Ptf1a*^CreER/WT^;*Kras*^LSL-G12D/WT^;*Ins1*^-/-^;*Ins2*^+/+^ mice were bred with *Ptf1a*^WT/WT^;*Ins1*^-/-^;*Ins2*^+/-^ mice to generate background-matched control mice (*Ptf1a*^CreER/WT^;*Kras*^LSL-G12D/WT^;*Ins1*^-/-^;*Ins2*^+/+^ mice), and experimental mice (*Ptf1a*^CreER/WT^;*Kras*^LSL-G12D/WT^;*Ins1*^-/-^;*Ins2*^+/-^ mice). The resulting litters were weaned to a high-fat diet with 60% fat (Research Diets D12492; Research Diets). At 4 weeks of age, mice were subcutaneously injected with tamoxifen (20 mg/mL in corn oil, Sigma-Aldrich) at 5 mg tamoxifen/40 g body mass for three consecutive days. At 57 weeks of age, mice were euthanized for histopathological analysis and single cell transcriptomics analysis. The *Kras*^LSL-G12D/WT^ mice (C57BL/6), *Ptf1a*^CreER/WT^ mice (C57BL/6), *Ins1*^-/-^ and *Ins2*^-/-^ mice (C57BL/6) were obtained as previously described (19).

### Glucose homeostasis and plasma amylase assessment

Mouse body weight, fasting glucose, and fasting insulin were measured after 4 hours of fasting in fresh, clean cages according to standard methods described previously (19). Body weight and fasting glucose were measured every 4 weeks and fasting insulin was measured every 3 months. At 52-weeks of age, glucose-stimulated (2g/kg) insulin secretion, intraperitoneal glucose tolerance test (2g/kg) and insulin tolerance test (0.75U/kg, Eli Lilly, USA) were performed as previously described (22, 24, 25). At 57-weeks of age, blood was collected from mice after 4 hours of fasting and blood samples were centrifuged at 4°C, at 10621rcf for 10 minutes to collect blood serum. Then the fasting amylase levels were measured with an amylase activity assay kit (MAK009-1KT, MilliporeSigma, MA, USA) according to the manufacturers’ instructions.

### Histological and morphological analysis

At 57-weeks of age, mice were euthanized and extracted pancreata were fixed in 4% paraformaldehyde for 24 hours followed by paraffin embedding. Seven-micron paraffin sections were collected from every embedded mouse pancreas for a total of 60 sections. Then evenly spaced sections were hematoxylin and eosin (H&E) stained and scanned as previously described (19, 28). Histopathological analysis was blindly conducted by Anni Zhang and verified by Janel Kopp and David Schaeffer. For each mouse pancreas, the PanIN area, ADM area and adipocytes area were analyzed on 3 H&E-stained sections, which were 140µm away from each other. The total pancreatic area, PanIN area and ADM area were measured as previously described (19). For the adipocyte occupied area, the pancreatic adipose tissue was selected and measured by Adobe Photoshop 2020 magic wand tool and measurement log function, respectively. The whole mouse pancreas area was selected by using Adobe Photoshop 2020 magnetic lasso tool and magic wand tool. Then the selected whole pancreas area was measured by the Adobe Photoshop 2020 measurement log function.

### Tissue dissociation and single-cell sorting

Six *Ptf1a*^CreER^;*Kras*^LSL-G12D^;*Ins1*^-/-^;*Ins2*^+/+^ mice pancreata and six *Ptf1a*^CreER^;*Kras*^LSL-G12D^;*Ins1*^-/-^;*Ins2*^+/-^ mice pancreata were dissociated for single cell RNA sequencing (scRNAseq) analysis (evenly divided by sex). Freshly dissected pancreata were washed with HBSS (Corning, 21-021-CV) and minced into ∼1mm pieces using sharp dissection scissors. Pieces were transferred into 15ml conical tubes and centrifuged at 720g for 2 minutes. The supernatant was discarded, and tissue was resuspended in 5l of ice-cold HBSS containing 0.4mg/ml collagenase P (Roche, #11213857001) and 10mg/ml DNase I (Roche, #11284932001). Tissue samples were incubated at 37°C water bath for 10-18 minutes and tubes were gently shaken with marbles. After the dissociation, 10ml HBSS + 5% FBS (ThermoFisher Scientific, #A3160701) was added to samples and sample were centrifuged at 720g for 2 minutes. Supernatants were discarded and sample were washed 3 times with 10ml HBSS + 5% FBS. Samples were filtered through a 100µM cell strainer then washed with HBSS + 5% FBS into a 50ml conical tube. Strainers were washed with 10ml HBSS + 5% FBS to collect the cells that remained on the strainer. Samples were centrifuged at 180g for 2 minutes to collect the dissociated cells. The resuspended samples were stained with 0.05μg/ml Hoechst 33342 (Invitrogen, #H3570) and 0.5μg/ml propidium iodide (Sigma, #P4864) and were sorted for live cells using BD LSR II Flow Cytometer.

### Single-cell transcriptomic data processing, quality control and analysis

The single cell libraries were prepared with the Chromium Single Cell 3’ Reagent Kits V3 (10X genomics, Pleasanton, CS, USA) according to the manufacturers’ instructions and the libraries were then sequenced on a Nextseq500 (Illumina). The Cell Ranger pipeline (10X genomics, CS, USA) was used to perform the demultiplexing (cellranger mkfastq, 10X genomics) and alignment (cellranger count, 10X genomics). The ambient RNA contamination was cleaned by R package SoupX (29). Cleaned gene-cell matrices were loaded into the R package Seurat 3.0 and filtered to remove cells with unique feature counts over 6000 or less than 200. The cells that had >20% mitochondrial gene counts or genes that were expressed by fewer than 3 cells were also removed. Using Seurat 3.2.1 (30, 31), the filtered gene-cell matrices from each mouse were integrated and clustered in uniform manifold approximation and projection (UMAP) space using default settings with resolution of 0.1. The typical marker genes (like *Prss2, Krt19*, and *Col1a1*) were used for identifying clusters. When the cluster identity could not be determined, Seurat 3.2.1 FindConservedMarkers function was used to find the top 50 genes that are conserved markers irrespective of the genotype. This gene list was then be uploaded to Enrichr and we used the suggested cell types by Enrichr to determine the identities for those clusters (32).

### Differential gene expression analysis, pathway enrichment analysis and visualization

A differential gene analysis was performed between two mouse genotypes (*Ptf1a*^CreER^; *Kras*^LSL-G12D^;*Ins1*^-/-^;*Ins2*^+/-^ vs *Ptf1a*^CreER^;*Kras*^LSL-G12D^;*Ins1*^-/-^;*Ins2*^+/+^) to identify upregulated and downregulated genes in each cell type using Seurat 3.2.1 FindMarkers function with default settings (30, 31). For the differentially expressed gene lists, the pathway enrichment analyses were performed by g:profiler (33) using the Reactome database (34) based on the Reimand *et al*. published protocol (35). The enriched pathways were visualized by R package pheatmap 1.0.12.

### Statistical analysis

Statistical parameters including the sample size n (number of animals), mean ± standard error of the mean (SEM), and statistical significance are reported in the figure legends and figures. GraphPad Prism 9.0.0 was used for performing the statistical analysis. Mixed-effects analysis was performed for glucose homeostasis assessments (Figure 1). When the samples passed a normality test, the two-tailed student’s t-test was performed; while non-parametric statistics (Mann-Whitney test) was run for data of non-normal distribution. GraphPad Prism 9.0.0 was used to generate and assess the linear regressions. Pearson correlation coefficients were computed for normally distributed data and nonparametric Spearman correlation was performed for non-normally distributed data. Non-parametric Wilcoxon rank sum test was used for differential gene expression analysis. p<0.05 was considered as significant and asterisks denote statistical significance level (*, p< 0.05; **, p<0.01; ***, p<0.001; ****, p<0.0001).

**Figure 1.**
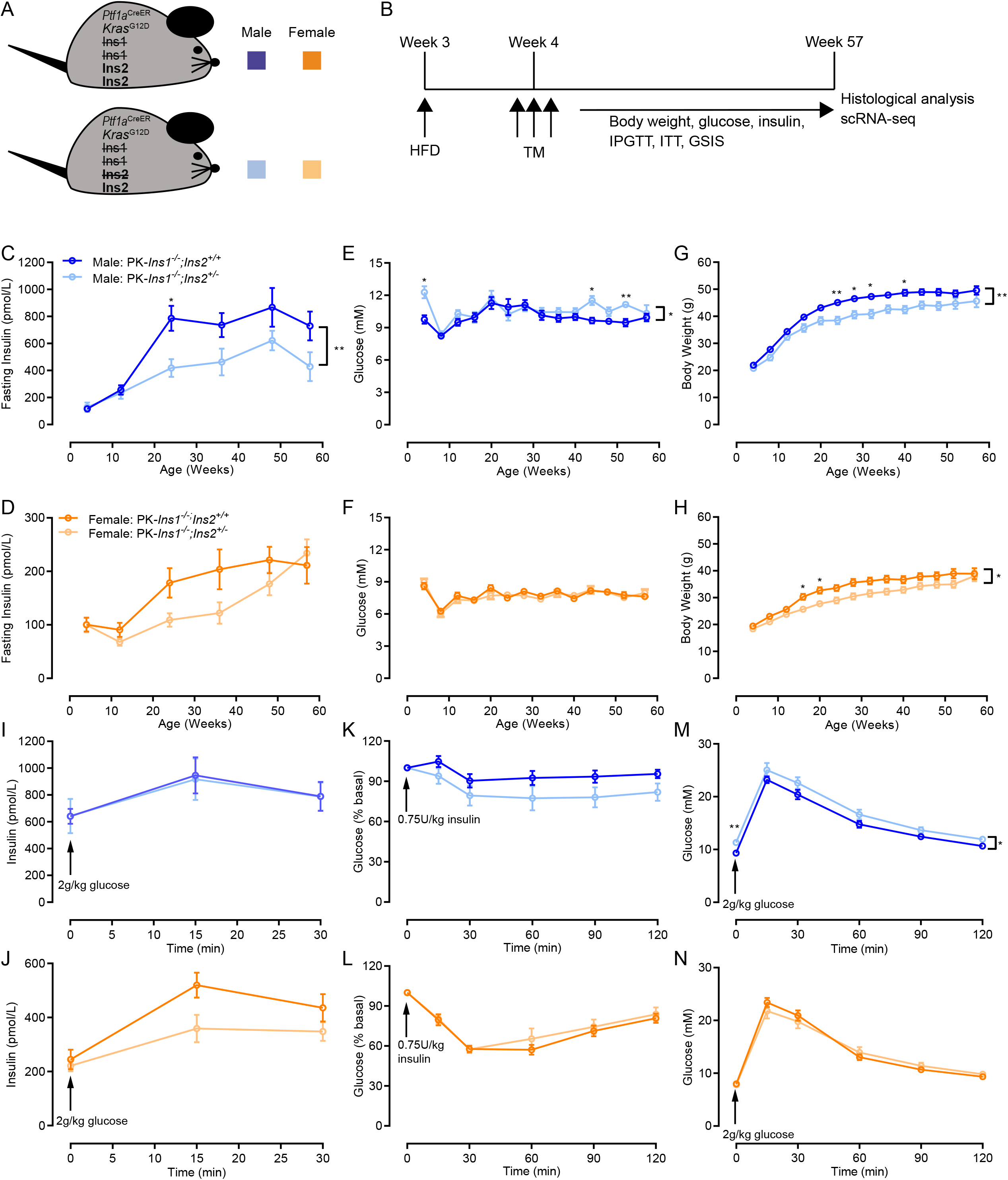
Mice with reduced insulin gene dosage have reduced fasting insulin levels and body weight. **(A)** Schematic describing a mouse model designed to test the role of insulin on HFD-accelerated PDAC initiation. On the background of *Ptf1a*^CreER^ -induced *Kras*^G12D^ pancreatic cancer model (PK), we compared experimental mice with 1 null allele of *Ins2* and control mice with 2 null alleles of *Ins2*, all in the absence of *Ins1* (*Ins1*^*-/-*^) to prevent compensation. **(B)** Three-week-old PK-*Ins1*^-/-^;*Ins2*^+/+^ control and PK-*Ins1*^-/-^;*Ins2*^+/-^ experimental mice were weaned to a high-fat diet (HFD) and were injected for 3 consecutive days with tamoxifen (TM) beginning at 4 weeks. After repeated physiological measures over the course of a year, the mice were then euthanized at 57 weeks of age for histological analysis and scRNA-seq. **(C-D)** Fasting insulin levels in male and female PK-*Ins1*^-/-^;*Ins2*^+/+^ and PK-*Ins1*^-/-^;*Ins2*^+/-^ mice measured over 1 year (n=18-29). **(E-F)** Fasting glucose levels in male and female PK-*Ins1*^-/-^;*Ins2*^+/+^ and PK-*Ins1*^-/-^;*Ins2*^+/-^ mice measured over 1 year (n=18-29). **(G-H)** Body weight in male and female PK-*Ins1*^-/-^;*Ins2*^+/+^ and PK-*Ins1*^-/-^;*Ins2*^+/-^ mice measured over 1 year (n=18-29). **(I-J)** Glucose stimulated insulin release in 52-week-old male and female mice (n=17-30). **(K-L)** Blood glucose response to intraperitoneal delivery of an insulin analog in 52-week-old male and female PK-*Ins1*^-/-^;*Ins2*^+/+^ and PK-*Ins1*^-/-^;*Ins2*^+/-^ mice (n=10-29). **(M-N)** Blood glucose response to intraperitoneal delivery of glucose in 52-week-old male and female PK-*Ins1*^-/-^;*Ins2*^+/+^ and PK-*Ins1*^-/-^;*Ins2*^+/-^ mice (n=16-29). *p<0.05 and **p<0.01. Values are shown as mean ± SEM.

## Results

### Effects of reduced *Ins2* gene dosage on hyperinsulinemia, obesity, and glucose homeostasis

We examined the effects of reduced *Ins2* gene dosage on PDAC development by crossing *Ptf1a*^CreER/WT^;*Kras*^LSL-G12D/WT^;*Ins1*^-/-^;*Ins2*^+/+^ mice with *Ptf1a*^WT/WT^;*Ins1*^-/-^;*Ins2*^+/-^ mice to activate mutant Kras expression in adult acinar cells and modulate insulin dosage (28, 36). By this breeding, we generated both *Ptf1a*^CreER^;*Kras*^LSL-G12D^;*Ins1*^-/-^;*Ins2*^+/+^ (PK-*Ins1*^-/-^;*Ins2*^+/+^) control mice and *Ptf1a*^CreER^;*Kras*^LSL-G12D^;*Ins1*^-/-^;*Ins2*^+/-^ (PK-*Ins1*^-/-^;*Ins2*^+/-^) experimental mice (Fig. 1A). Recombination and expression of the *Kras*^LSL-G12D^ allele was induced by injecting mice with tamoxifen at 4 weeks of age. To stimulate high insulin production, the mice were fed a HFD after weaning (Fig. 1B).

In contrast to our previous study, which had more limited physiological data, especially in males (19), body weight and fasting glucose levels were monitored every 4 weeks and fasting insulin levels were measured every 3 months until euthanasia in both male and female mice (Fig. 1B). As expected from our previous studies using mice with reduced insulin gene dosage (24, 25, 37, 38), PK-*Ins1*^-/-^;*Ins2*^+/-^ mice had lower fasting insulin levels than PK-*Ins1*^-/-^;*Ins2*^+/+^ mice for both male and female mice (Fig. 1C-D). However, as previously reported (25), this effect was transient in females over this time period (Fig. 1D). Mice with reduced fasting insulin levels exhibited reduced weight gain in the context of HFD, without consistently affecting glucose homeostasis (Fig. 1E-H), consistent our previous reports (22, 24, 25, 38). When mice were 52-weeks-old, glucose-stimulated insulin secretion tests, insulin tolerance tests, and glucose tolerance tests were conducted to examine glucose homeostasis more closely (Fig. 1I-N). At this age, PK-*Ins1*^-/-^;*Ins2*^+/+^ and PK-*Ins1*^-/-^;*Ins2*^+/-^ mice secreted statistically similar levels of insulin in response to intraperitoneal delivery of glucose, although experimental female mice showed a clear trend towards reduced glucose responsiveness (Fig. 1I-J). Male mice were generally more insulin resistant than female mice (Fig. 1K-L). However, there were no significant differences in insulin sensitivity between the genotypes, even though males with reduced insulin production appeared to be slightly more insulin sensitive. Glucose tolerance was generally similar, although male mice with reduced insulin were slightly, but significantly, more intolerant to glucose than *Ins2*^+/+^ littermate controls (Fig. 1M-N). We also examined any potential differences in exocrine physiology by monitoring serum amylase levels (39), but found no differences between the genotypes (Fig. S1A-B).

In sum, the limited systemic physiological differences between the experimental PK-*Ins1*^-/-^;*Ins2*^+/-^ and control PK-*Ins1*^-/-^;*Ins2*^+/+^ mice offered an opportunity to examine the effects of reduced insulin production on PanIN formation in the absence of major changes in glucose homeostasis in both sexes.

### Effects of modestly reduced insulin on PanIN initiation

Mice were euthanized at 57 weeks of age for histological analysis of the percent of total pancreatic area occupied by PanIN and tumor or for scRNAseq analysis (see below). We detected ductal lesions with histologic characteristics of low-grade PanIN (Fig. 2A-B). Similar to our previous study, the pancreatic area covered by PanIN and tumor in the PK-*Ins1*^-/-^;*Ins2*^+/+^ control mice was approximately twice that of the PK-*Ins1*^-/-^;*Ins2*^+/-^ experimental mice with reduced insulin levels (1.34% ± 3.41% vs 0.36% ± 0.26%, respectively for males and 3.54% ± 8.03% vs 1.58% ± 2.53%, respectively for females) (Fig. 2C-D). However, unlike our previous study with varying alleles of *Ins1* in an *Ins2*-null background (19); we observed far fewer PanIN lesions in both genotypes and the difference in PanIN area was not statistically significant. This is perhaps related to the overall reduction in non-fat pancreatic area in the *Ins1*-null compared to the *Ins2*-null background (see below). Similar to our previous study, only one male and one female mouse developed tumors and both of them were from PK-*Ins1*^-/-^;*Ins2*^+/+^ genotype (Fig. 2C-D, filled circles). Next, we examined the correlations between PanIN plus tumor area and fasting insulin levels, glucose levels, and body weight in individual mice measured at 57 weeks of age, by pooled measurements (black) or within each group (colored) (Fig. 2E-J). Relatively modest positive correlations between fasting insulin levels and PanIN plus tumor area were only significant in female mice (Fig. 2F). Female mice also had a significant, but even more modest, correlation between body weight and PanIN plus tumor area (Fig. 2J). There was no positive correlation between glucose and PanIN plus tumor area in either sex (Fig. 2G-H), consistent with our previous findings (19). Together, these data add support to our previous observations suggesting that hyperinsulinemia promotes PanIN development (19).

**Figure 2.**
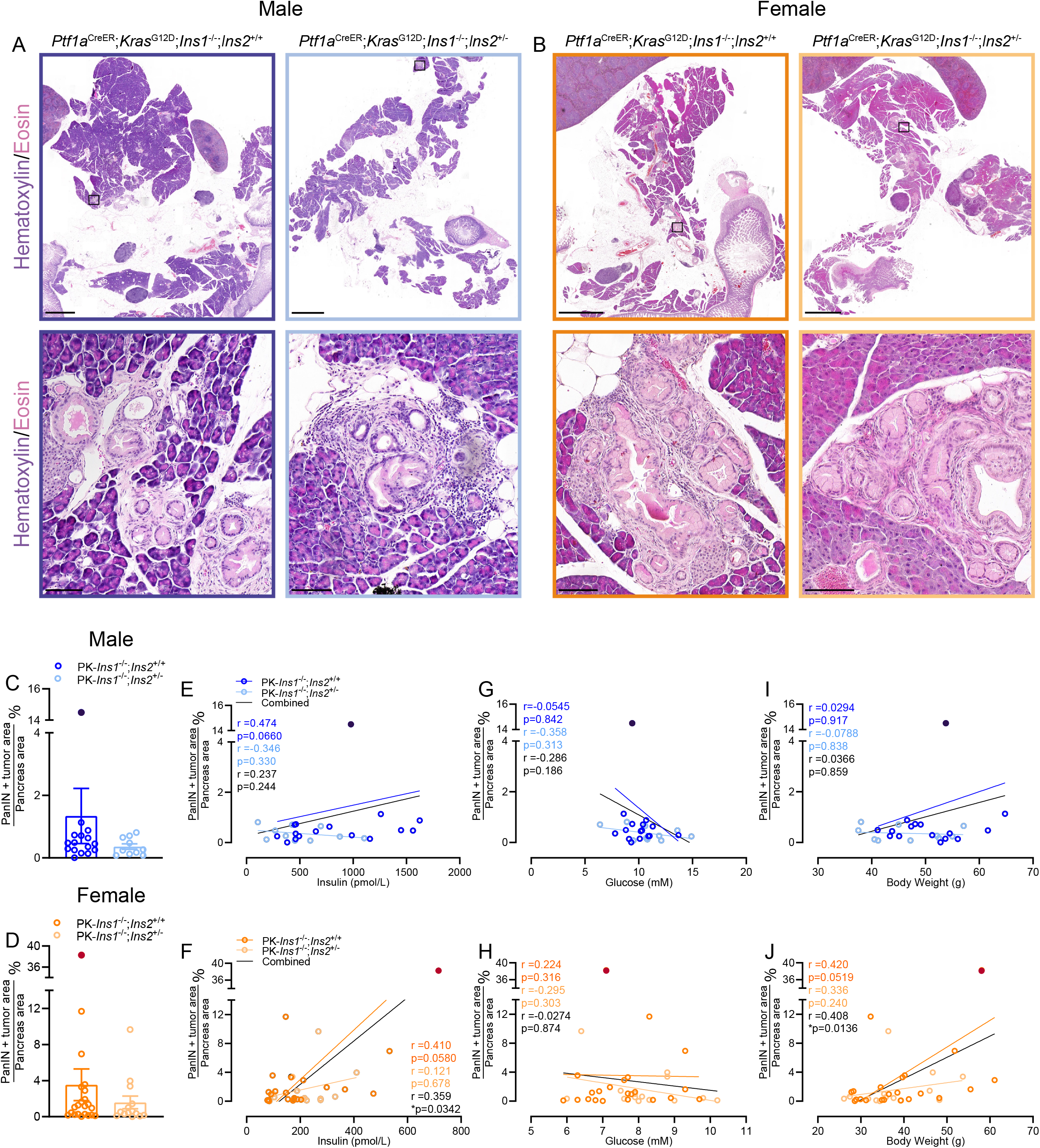
Effects of reduced hyperinsulinemia on pancreatic cancer initiation. **(A-B)** Representative whole section (*top*) and high-magnification (*bottom*) images of PK-*Ins1*^-/-^;*Ins2*^+/+^ and PK-*Ins1*^-/-^;*Ins2*^+/-^ male and female pancreata stained with hematoxylin and eosin. Scale bars: 2 mm (top) and 0.1mm (bottom). **(C-D)** Quantification of percent of total pancreatic area occupied by PanINs and tumor in male (blue colors) and female (orange colors) PK-*Ins1*^-/-^; *Ins2*^+/+^ and PK-*Ins1*^-/-^;*Ins2*^+/-^ mice (n= 14-22) (dark blue and dark orange dots denote mice that developed tumors). Correlations of composite PanINs plus tumor area with fasting insulin levels **(E-F)**, with fasting glucose levels **(G-H)**, or with body weight **(I-J)** in male and female PK-*Ins1*^-/-^;*Ins2*^+/+^ and PK-*Ins1*^-/-^;*Ins2*^+/-^ mice (n = 10-22).

### Acinar ductal metaplasia and adipocyte area in mice with reduced hyperinsulinemia

Next, we measured the percent of total pancreatic area covered by ADM. ADM is the histological evidence of normal acinar cells changing into ductal-like cells with ductal cell morphology and it can be induced by pancreatitis and during PanIN development (40). We detected ADM in both male and female mice for each genotype (Fig. 3A-B). Similar to percent of PanIN area, PK-*Ins1*^-/-^;*Ins2*^+/+^ mice had two times the amount of ADM area as PK-*Ins1*^-/-^;*Ins2*^+/-^ mice (1.92% ± 3.33% vs 0.96% ± 0.99%, respectively for males and 9.53% ± 13.00% vs 4.45% ± 6.29%, respectively for females), but the difference was not statistically significant (Fig. 3C-D). We also examined the correlations between percent ADM area and fasting insulin, fasting glucose, and body weight, for each genotype (orange or blue) or both together (black) (Fig. 3E-J). Although there was a trend for the ADM area to correlate with fasting insulin in both sexes, only when the genotypes were combined to increase power did the percent ADM area significantly correlate with fasting insulin in males (Fig. 3E) or body weight in females (Fig. 3J).

**Figure 3.**
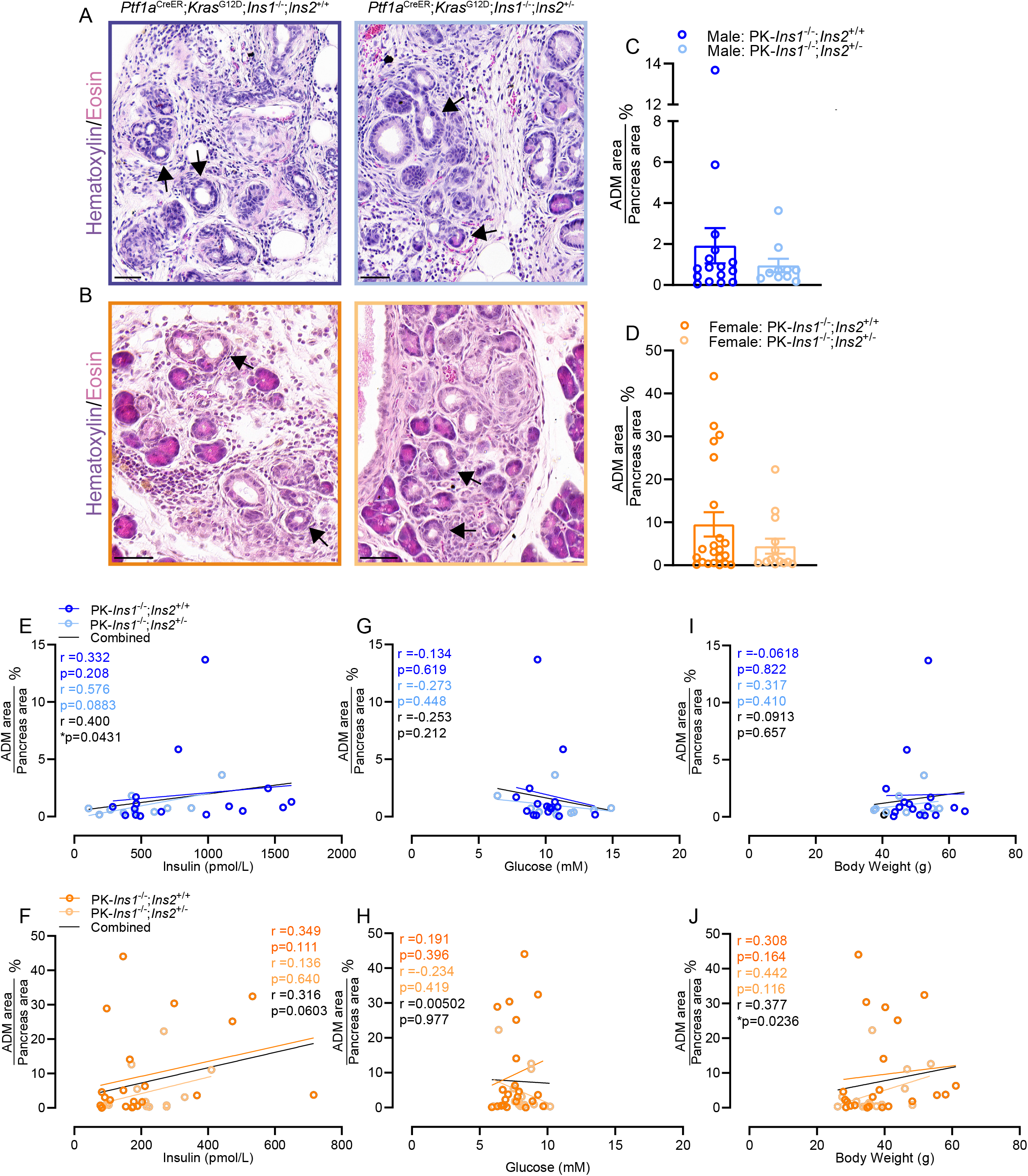
Altered acinar ductal metaplasia (ADM) in mice with reduced hyperinsulinemia. **(A-B)** Representative high-magnification ADM (arrowheads) images of male **(A)** and female **(B)** PK-*Ins1*^-/-^;*Ins2*^+/+^ (*left*) and PK-*Ins1*^-/-^;*Ins2*^+/-^ (*right*) mouse pancreas sections stained with H&E. **(C-D)** Quantification of percent of total pancreatic area occupied by ADM in male and female mice of each genotype (n= 10-22). Correlations of composite ADM area with fasting insulin levels **(E-F)**, fasting glucose **(G-H)**, or body weight **(I-J)** in male **(E, G, I)** and female **(F, H, I)** PK-*Ins1*^-/-^;*Ins2*^+/+^ and PK-*Ins1*^-/-^;*Ins2*^+/-^ mice (n = 10-16). Values are shown as mean ± SEM. Scale bars: 0.05mm.

One surprising observation from our histological analyses was the significant amount of pancreatic area that had been replaced by adipocytes in our PK-*Ins1*^-/-^ mouse model. This is not a phenomenon that we had previously observed in our PK-*Ins2*^-/-^ models (19). As the representative histological figures show (Fig. 4A-B), we often observed pancreatic lobules with a few residual normal acinar, ductal, or endocrine cells left amongst large numbers of adipocytes. The percent of pancreatic area replaced by adipocytes was not significantly different between PK-*Ins1*^-/-^;*Ins2*^+/+^ and PK-*Ins1*^-/-^;*Ins2*^+/-^ mice (Fig. 4C-D), suggesting this phenotype was possibly associated with loss of *Ins1* gene specifically. The fatty replacement affected the overall parenchyma area, as we found compared to PK-*Ins2*^-/-^ female mice, PK-*Ins1*^-/-^ female mice had significantly less pancreatic area (PK-*Ins2*^-/-^ male mice could not be assessed)(Fig. S1C). It is possible that this fatty replacement could have affected the overall number of PanIN lesions, because of a relative lack of Ptf1a-positive acinar cells. There was a significant correlation between the percent of adipocyte area and fasting insulin for female, but not male mice (Fig. 4E-F). We observed no correlation between percent of adipocyte area and fasting glucose levels for either sex (Fig. 4G-H). However, as expected (25), the percent adipocyte area did correlate with body weight in both sexes (Fig. 4I-J). The underlying cause of fatty replacement in the PK-*Ins1*^-/-^ mice is unknown, but it could potentially influence the accumulation of PanIN lesions. We have previously observed this fatty replacement of normal parenchyma in another *Ins1*^-/-^ colony of mice (unpublished observations), therefore, we do not believe this phenomena is solely related to the exposure of mice to tamoxifen or the influence of the PK mutant alleles.

**Figure 4.**
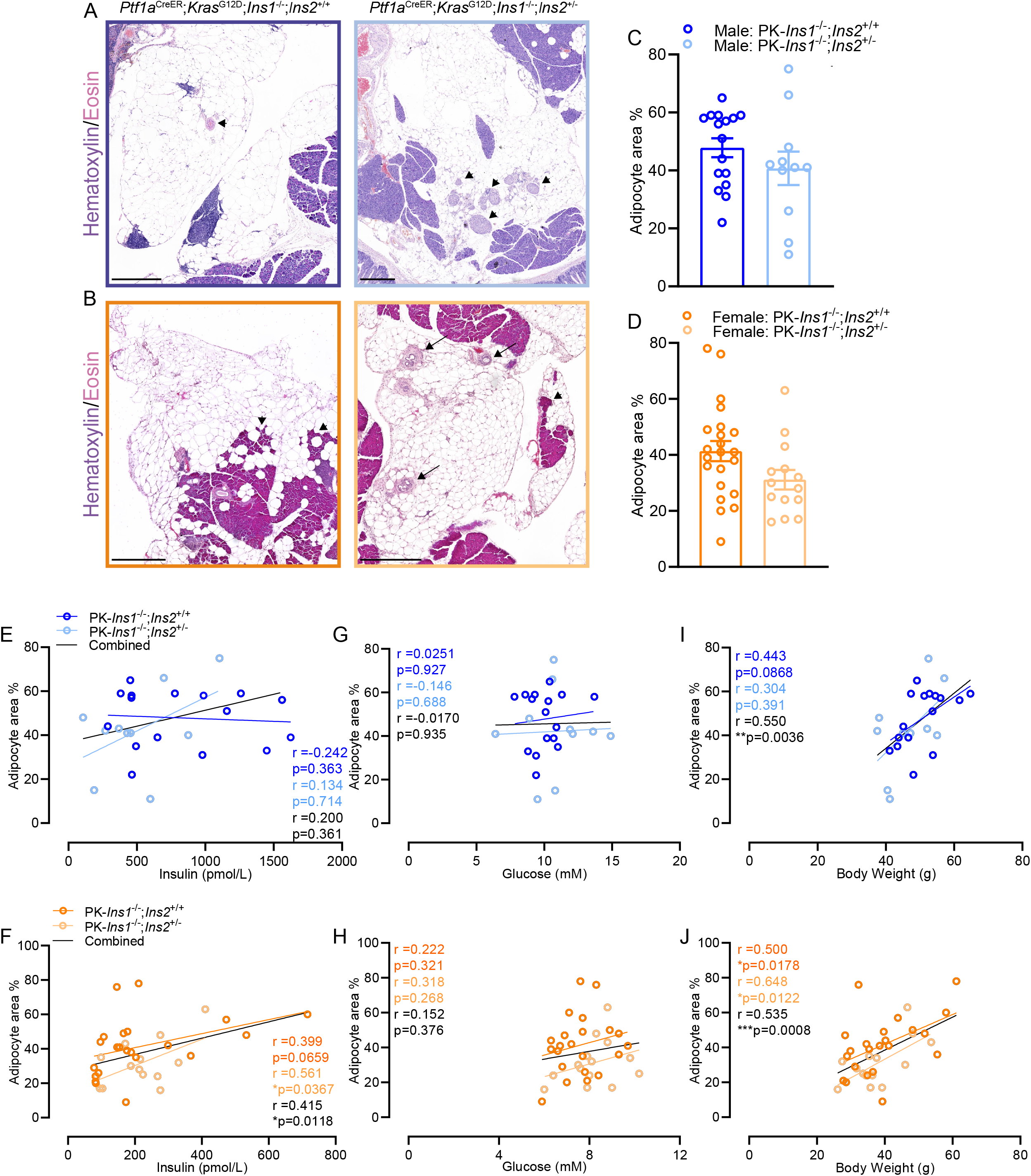
Altered pancreatic adipocyte area in mice with reduced hyperinsulinemia. **(A-B)** Representative high-magnification image of apparent adipocyte replacement of pancreas area in male **(A)** and female **(B)** PK-*Ins1*^-/-^;*Ins2*^+/+^ (left) and PK-*Ins1*^-/-^;*Ins2*^+/-^ (right) mice stained with H&E. Black arrows point to residual ducts. Arrowheads point to remaining islets, acinar cells, and blood vessel. **(C-D)** Quantification of percent of total pancreatic area occupied by adipocytes in male and female mice of each genotype (n= 11-22). Correlations of adipocyte area with fasting insulin levels **(E-F)**, fasting glucose levels **(G-H)**, or body weight **(I-J)** for male **(E, G, I)** and female **(F, H, J)** PK-*Ins1*^-/-^;*Ins2*^+/+^ (dark colors) and PK-*Ins1*^-/-^;*Ins2*^+/-^ (light colors) mice. (n = 11-22). Values are shown as mean ± SEM. Scale bars: 0.5mm.

### Single-cell transcriptomics reveals effects of hyperinsulinemia on cell type-specific gene expression

To investigate the molecular effects of hyperinsulinemia in the context of PDAC initiation in an unbiased and cell type-specific manner, we undertook scRNAseq. Only a few studies have successfully conducted scRNAseq in the pancreas to date for studying pancreatic cancer (41-43). At 57 weeks of age, we collected pancreata from 6 PK-*Ins1*^-/-^;*Ins2*^+/+^ control mice and 6 PK-*Ins1*^-/-^;*Ins2*^+/-^ experimental mice, dispersed them into single cells, and FACS purified live cells for sending to single-cell RNA sequencing (Fig. 5A). In total, 49,835 single cells passed quality control tests and were clustered into 15 clusters (Fig. 5B-C). These cell clusters were assigned cellular identities based on the expression of known markers (Fig. 5D). We were able to identify acinar cells, ductal cells, and fibroblasts. The majority of cells that survived dispersion and passed transcriptomics quality controls were immune cells including: T cells, T regulatory cells (Treg), B cells, natural killer (NK) cells, macrophages (both M1 and M2 macrophages), monocytes, dendritic cells, and mast cells (Fig. 5B-D, and Fig. S2D). We also classified a separate cluster of proliferating cells marked by high expression of *Mki67*. This proliferating cell cluster also included multiple immune cell types, such as T cells, B cells, and NK cells, as well as epithelial cells (Fig. S2A-C). With the exception of twice as many NK cells in mice with reduced insulin production, there were no significant differences in the numbers of cells per cluster between the genotypes (Fig. 5C). Analysis of cell type specific markers showed that cell identities were generally comparable between genotypes (Fig. 5D).

**Figure 5.**
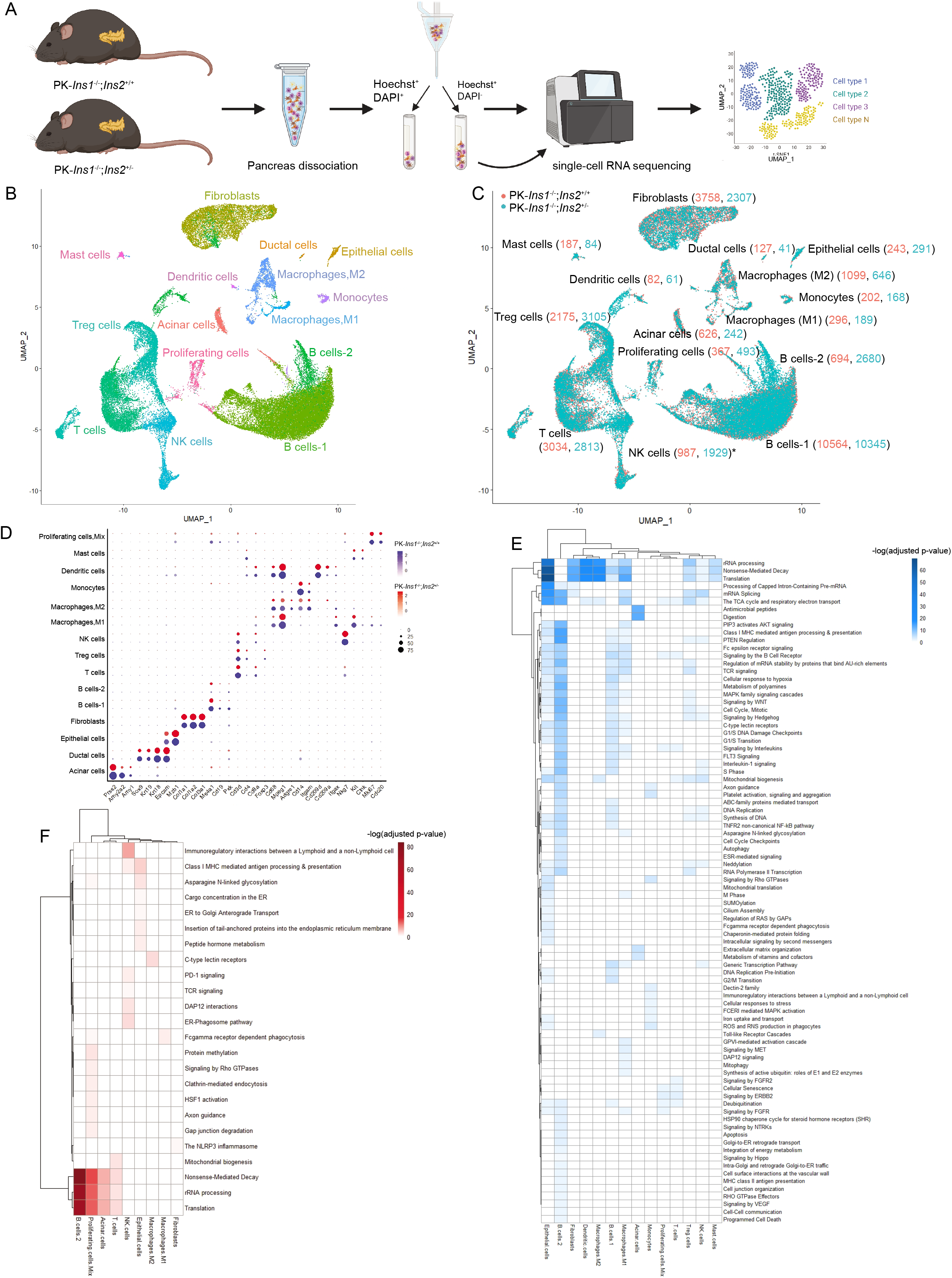
scRNAseq analysis reveals effects of hyperinsulinemia on cell type-specific gene expression. **(A)** Schematic describing the single-cell transcriptomics experimental design and analysis. Six mouse pancreata from each genotype were dissociated to single cells and the samples were sorted for Hoechst-positive and DAPI-negative live cells. The live single cells were sequenced and clustered in uniform manifold approximation and projection (UMAP) space. **(B)** Unsupervised clustering of cells from 6 PK-*Ins1*^-/-^;*Ins2*^+/+^ and 6 PK-*Ins1*^-/-^;*Ins2*^+/-^ mice pancreata, represented as an UMAP plot. **(C)** Numbers of cells from PK-*Ins1*^-/-^;*Ins2*^+/+^ (green) and PK-*Ins1*^-/-^;*Ins2*^+/-^ (orange) mice for each cell type. The asterisk indicates a significant difference in the number of NK cells between the genotypes. **(D)** Dot plot showing selected cell type-specific markers for identifying the cell type for each cluster. The size of dots represents the fraction of cells expressing the markers. The PK-*Ins1*^-/-^; *Ins2*^+/+^ mouse data are shown in blue and PK-*Ins1*^-/-^;*Ins2*^+/-^ mouse data are shown in red. The intensity of color indicates the average expression of marker genes for each cell type. **(E)** Heatmap showing Reactome pathways that are downregulated in PK-*Ins1*^-/-^;*Ins2*^+/-^ mice when compared to PK-*Ins1*^-/-^;*Ins2*^+/+^ mice for each cell type. The intensity of color indicates the negative log_10_ of adjusted p value. **(F)** Heatmap of pathways that are upregulated in PK-*Ins1*^-/-^;*Ins2*^+/-^ mice when compared to PK-*Ins1*^-/-^;*Ins2*^+/+^ mice for each cell type. The intensity of color indicates the negative log_10_ of adjusted p value. *p<0.05.

After confirming the cell identities, we generated a list of genes that were differentially expressed between genotypes for each cell type (Supplemental table1). To have a more comprehensive understanding of the function of these differentially expressed genes, we performed pathway enrichment analysis using Reactome and found the pathways that were down-(Fig. 5E) and up-regulated (Fig. 5F) in PK-*Ins1*^-/-^;*Ins2*^+/-^ experimental mice compared to PK-*Ins1*^-/-^;*Ins2*^+/+^ control mice. B cells-1, B cells-2, epithelial cells and M1 macrophages were the cell types that had the most altered pathways (Fig. 5E-F). Specifically, the pathways that were most significantly altered were rRNA processing, nonsense-meditated decay, and translation pathways and they were also the pathways that were consistently altered across multiple cell types (Fig. 5E-F). Interestingly, they were down-regulated in epithelial cells, fibroblasts, dendritic cells, macrophages, B cells-1, Treg, NK cells and mast cells of mice with reduced insulin. However, they were up-regulated in B cells-2, proliferating cells, and acinar cells of mice with reduced insulin (Fig. 5E-F). We also found pathways that were only altered in one cell type. For instance, antimicrobial peptides and digestion pathways were down-regulated in acinar cells, while PD-1 signaling and TCR signaling were up-regulated in NK cells from PK-*Ins1*^-/-^;*Ins2*^+/-^ experimental mice (Fig. 5E-F). *Reg3a, Reg3b, Reg3d*, and *Reg3g* genes, which are known to be induced by inflammation and may have anti-microbial roles (44, 45), were also significantly downregulated in acinar cells from mice with reduced insulin production (Fig. S2E). Reg proteins have been demonstrated to be able to promote pancreatic carcinogenesis (46-48).

Somewhat expectedly, pathways involved in insulin signaling, like “PIP3 activates AKT signaling”, “PTEN regulation”, and “MAPK family signaling cascades,” were downregulated in mice with reduced insulin production. They were downregulated in several cell types, but most clearly in B cells and M1 macrophages (Fig. 5E-F). There was also a downregulation of genes involved in cell cycle pathways in epithelial cells, B cells, macrophages, Treg cells, and NK cells from mice with reduced insulin production (Fig. 5E). Altogether, the pathway enrichment analysis suggested that genes involved in translation were most significantly and consistently altered by hyperinsulinemia. This suggests that hyperinsulinemia might indirectly affect PanIN development through regulating the immune cells in the PanIN microenvironment.

## Discussion

The goal of this study was to investigate the effects of reduced *Ins2* gene dosage on HFD-induced hyperinsulinemia, PanIN initiation, and cell type specific gene expression in the context of acinar-cell-specific expression of mutant Kras. The results of the present study extend our previous findings (19), which implicate hyperinsulinemia as a causal factor in pancreatic cancer initiation and provide the first molecular insights into the cell-specific mechanisms involved.

Despite the strong epidemiological link between hyperinsulinemia and pancreatic cancer, the specific reduction of insulin is required to formally test the hypothesis that insulin plays a causal role. Our previous study was the first to demonstrate that endogenous hyperinsulinemia contributes to cancer development using mice with reduced dosage of *Ins1* in a *Ins2*-null genetic background (19). Unfortunately, in that study male PK-*Ins1*^+/-^;*Ins2*^-/-^ mice developed hyperglycemia at very young age because of insufficient endogenous insulin production, which limited our strongest conclusions to only female mice (19). In the present study, we were eager to extend our observations to both sexes and indeed, we found that male PK-*Ins1*^-/-^;*Ins2*^+/-^ mice were able to maintain glucose homeostasis and be studied long-term. This is consistent with previous studies showing that limiting *Ins2* gene dosage prevented hyperinsulinemia without affecting glucose homeostasis (24). Our data show that reduced *Ins2* gene dosage led to a moderate reduction in fasting insulin levels without affecting glucose homeostasis, in both male and female mice. It should be noted that circulating insulin levels in female mice, even with both *Ins2* alleles, is only ∼25% of that seen in male mice. We also noted in female mice that insulin levels were not different at 1 year of age between genotypes, mirroring the transient compensation we have previously observed in *Ins1*-null model (25, 38). Collectively, these observations illustrate that a reduction in *Ins2* gene dosage results in a relatively mild manipulation of circulating insulin in the first year of life. Because *Ins2* is the ancestral gene and contributes to ∼2/3 of secreted insulin (21,23), fasting insulin levels were still relatively high for our PK-*Ins1*^*-/-*^ mouse model compared to the previously studied PK-*Ins2*^*-/-*^ mouse model.

In the present study, we conducted extensive histological analysis of PanIN, ADM, and fatty replacement. Interestingly, in the present study only about 1-4% pancreas was occupied by PanIN lesions for our PK-*Ins1*^*-/-*^ mouse model, compared with the 15-30% of pancreas that was occupied by PanIN lesions for our previous PK-*Ins2*^*-/-*^ mouse model (19). Another major histological difference between this study and our previous one, was the observation of a significant amount of fatty replacement in our PK-*Ins1*^-/-^ mouse model. Approximately 30-50% of the pancreas was replaced by adipocytes. Pancreatitis can induce acinar cell necrosis or apoptosis, which is subsequently replaced by adipocytes (49, 50). HFD-induced obesity can also cause fat accumulation in the exocrine parenchyma (49, 51), but we did not observe this extent of fatty replacement with the same diet in our previous model (19). Together it seems that the combined effects of Kras-associated inflammation, HFD, and the *Ins1*^-/-^ genetic background may have resulted in fat displacing ∼2/3 of normal pancreatic parenchyma in our PK-*Ins1*^*-/-*^ mouse model. Therefore, the loss of acinar cells may explain why fewer and more variable numbers of PanIN lesions developed in our PK-*Ins1*^*-/-*^ mouse model. Nevertheless, circulating insulin was still significantly correlated with PanIN plus tumour area in female mice, confirming our previous report with another insulin gene dosage configuration. Our histopathological analysis also showed that PK-*Ins1*^-/-^;*Ins2*^+/-^ mice had a ∼50% reduction in percent PanIN area compared with PK-*Ins1*^-/-^;*Ins2*^+/+^ mice, which is similar to our previous findings. Although the pairwise comparison between genotypes did not reach statistical significance in the present study (19), our findings still support a role for hyperinsulinemia promoting PanIN initiation from acinar cells sustaining mutations in the oncogene *Kras*.

Pancreatic cancer is one of the most stroma-rich solid tumor types and immune cells in the microenvironment play important roles at both early and late stages of PDAC (52-55). In our single-cell transcriptomics analysis, the primary cell types analyzed were immune cells, including: T cells, B cells, macrophages, NK cells, and dendritic cells. As expected, the major signaling pathways downstream of insulin (e.g. MAPK-ERK, PI3K-AKT, cell cycle and translation pathways) were downregulated in immune cells from mice with reduced insulin production. MAPK-ERK and PI3K-AKT signaling pathways and their downstream signaling cascades are well established regulators of multiple immune cell types (12, 56, 57).

Infiltrating immune cells can play different roles in PanIN initiation. For instance, Zhang *et al*. found CD4^+^ T cells could repress antineoplastic function of CD8^+^ T cells as the depletion of CD4^+^ T cells reduced PanIN lesions in the presence of CD8^+^ T cells (55). High tumor infiltration of Tregs is usually considered as an unfavorable prognostic factor for PDAC (52, 58); however, a recent study found that ablation of Tregs during PanIN initiation accelerated pancreatic carcinogenesis as it caused a loss of tumor-restraining fibroblasts (59). The depletion of Tregs also increased myeloid cells recruitment which were shown to be required for establishing an immunosuppressive microenvironment during PanIN development (12, 59, 60). Our data suggested that there were more Tregs and less myeloid cells (macrophages and monocytes) from mice with reduced insulin production. This composition of immune cells may hinder PanIN initiation and contribute to the reduced PanIN area in PK-*Ins1*^-/-^;*Ins2*^+/-^ mice. Additionally, macrophages also affect PDAC development. Macrophages can be either proinflammatory (M1) with anti-tumor properties, or anti-inflammatory (M2) with tumor-promoting actions (52, 54, 58), and they are recruited to pancreas at early stages of tumorigenesis (53, 61, 62). Previous studies showed that M2 macrophages were the predominate phenotype of macrophages detected around PanIN lesions (51, 63), and PanIN cells could produce cytokines like interleukin-13 to drive M2 polarization (64). M2 macrophages could contribute PanIN development through mediating fibrogenesis, angiogenesis and creating an immunosuppressive environment and the depletion of M2 macrophages significantly attenuated PanIN progression (51, 62). Interestingly, compared to PK-*Ins1*^-/-^;*Ins2*^+/+^ mice, the number of macrophages with an M2-like transcriptional profile was reduced ∼50% in PK-*Ins1*^-/-^;*Ins2*^+/-^ mice. In contrast, we observed a significant increase in the number of NK cells in our PK-*Ins1*^-/-^;*Ins2*^+/-^ mice. The deficiency in NK cells and natural killer T cells infiltration has been demonstrated to promote PanIN initiation (65, 66). Together, these observations suggested that the immune cells composition in mice with reduced insulin might have anti-tumorigenesis effects. In our mouse models, B cells had the greatest number of pathways altered by hyperinsulinemia.

The role of B cells in PDAC appears to be complex according to the few studies in which it has been investigated. Some studies demonstrated that B cells secreted interleukin-35 and promoted tumor progression, while Spear *et al*. showed that B cells were proinflammatory and limited PDAC development (67-69). Future studies are therefore required to better understand how hyperinsulinemia effects on B cells and how B cells contribute to PanIN development. Overall, our scRNAseq analysis suggested hyperinsulinemia might contribute to PanIN development through inducing PanIN-promoting properties of immune cells.

One of the limitations of our single-cell transcriptomics analysis was that relatively few acinar cells survived the pancreas dispersion and were gated as healthy prior to transcriptomic analysis. This was somewhat expected given the fragility of acinar cells and meant that we had less cell-level power to detect differences in gene expression and that we may not have been assessing gene expression in an exceptionally robust sub-grouping of acinar cells. Nevertheless, within the limited data, we observed a significant downregulation of *Reg3a, Reg3b, Reg3d*, and *Reg3g* in acinar cells from PK-*Ins1*^-/-^;*Ins2*^+/-^ mice. Reg proteins have been shown to promote pancreatic carcinogenesis, especially inflammation-linked pancreatic carcinogenesis (46-48). Inflammation in pancreas can cause ADM and accelerate PanIN progression (36, 70) and therefore, the decrease of *Reg* transcripts in PK-*Ins1*^-/-^;*Ins2*^+/-^ mice is consistent with the reduction of PanINs and inflammation in those mice. Future studies may seek to directly manipulate Reg proteins in the context of hyperinsulinemia and pancreatic cancer.

In future studies, it will be important to characterize the specific downstream changes for each immune cell type at the protein level. It will also be important to manipulate insulin signaling components, including the insulin receptor, in acinar cells, immune cells and other components of the PanIN microenvironment to determine which cells are predominately being affected by changes in insulin.

## Conclusions

The present study showed that mice with reduced hyperinsulinemia trended to have less pancreatic area covered by PanIN and ADM, consistent with our previous data demonstrating that hyperinsulinemia can contribute causally to PanIN development. The scRNA-seq analysis demonstrated that hyperinsulinemia affected the immune cell composition in the PanIN microenvironment and altered cellular pathways involved in or targets of insulin signaling, such as the MAPK-ERK pathways and protein translation. Gene expression changes in the PanIN immune microenvironment in the mice with reduced insulin production would be predicted to result in fewer PanIN lesions. Our study represents an important first step in understanding the molecular effects of hyperinsulinemia on all the cell types present in the context of early-stage pancreatic cancer.

### List of abbreviations

PanIN: Pancreatic intraepithelial neoplasia
ADM: Acinar-to-ductal metaplasia
PDAC: Pancreatic ductal adenocarcinoma
HFD: high fat diet
H&E: hematoxylin and eosin
scRNAseq: single cell RNA sequencing
UMAP: uniform manifold approximation and projection
SEM: standard error of the mean

## Declarations

### Ethics approval and consent to participate

University of British Columbia Animal Care Committee in accordance with Canadian Council for Animal Care guidelines approved all animal experiments.

## Consent for publication

Not applicable

## Availability of data and materials

All data generated and analyzed in this study is included within the article, additional files or is available from the corresponding author on request.

## Author’s contributions

A.M.Y.Z designed, managed, and conducted the project including acquiring, analyzing, and interpreting all data and wrote the manuscript (in vivo animal experiments, genotyping, histology, and scRNAseq). T.J.J.dW performed *in vivo* animal experiments and genotyping. S.F, Y.B.Z and, S.W provided advice on scRNAseq analysis. X.H and H.L provided help on *in vivo* animal experiments. JLK and JDJ supervised the project, obtained funding, interpreted the data, and edited the manuscript.

## Acknowledgments

The authors thank Audrey Hendley for the pancreas dissociation protocol and members from Johnson and Kopp labs for discussions.

## Supplemental Figure Legends

**Figure S1. Amylase activity and pancreatic area of PK-*Ins1***^**-/-**^**;*Ins2***^**+/+**^ **and PK-*Ins1***^**-/-**^**;*Ins2***^**+/-**^ **mice. (A-B)** The amylase activity in male **(A)** and female mice **(B)** for each genotype. **(C)** The total pancreatic area for mice in an *Ins2*-null background or in an *Ins1*-null background. ****p<0.0001. Values are shown as mean ± SEM.

**Figure S2. Proliferating cell cluster contains multiple cell types. (A)** Unsupervised sub-clustering of the cluster containing proliferating cells, represented as an UMAP plot. The proliferating cell cluster contains proliferating T cells, B cells, Naïve B cells, NK cells and epithelial cells. **(B)** Numbers of cells from PK-*Ins1*^-/-^;*Ins2*^+/+^ (green) and PK-*Ins1*^-/-^;*Ins2*^+/-^ (orange) mice for each cell type. **(C)** Violin plot showing the expression level of selected cell type-specific markers for identified cell types within the proliferating cell cluster. **(D)** Expression level of the typical markers for identifying M1 macrophages and M2 macrophages for each genotype. **(E)** The differential expression of *Reg3a, Reg3b, Reg3d*, and *Reg3g* genes in acinar cells between PK-*Ins1*^-/-^;*Ins2*^+/+^ and PK-*Ins1*^-/-^;*Ins2*^+/-^ mice.

## References

1. Ilic M, Ilic I. Epidemiology of pancreatic cancer. World J Gastroenterol. 2016;22(44):9694–705.

2. Renehan AG, Tyson M, Egger M, Heller RF, Zwahlen M. Body-mass index and incidence of cancer: a systematic review and meta-analysis of prospective observational studies. Lancet. 2008;371(9612):569–78.

3. Hart AR, Kennedy H, Harvey I. Pancreatic cancer: a review of the evidence on causation. Clin Gastroenterol Hepatol. 2008;6(3):275–82.

4. Koorstra JB, Hustinx SR, Offerhaus GJ, Maitra A. Pancreatic carcinogenesis. Pancreatology. 2008;8(2):110–25.

5. Permuth-Wey J, Egan KM. Family history is a significant risk factor for pancreatic cancer: results from a systematic review and meta-analysis. Fam Cancer. 2009;8(2):109–17.

6. Lowenfels AB, Maisonneuve P. Risk factors for pancreatic cancer. J Cell Biochem. 2005;95(4):649–56.

7. Lauby-Secretan B, Scoccianti C, Loomis D, Grosse Y, Bianchini F, Straif K, et al. Body Fatness and Cancer--Viewpoint of the IARC Working Group. N Engl J Med. 2016;375(8):794–8.

8. Gallagher EJ, LeRoith D. Hyperinsulinaemia in cancer. Nat Rev Cancer. 2020;20(11):629–44.

9. Gallagher EJ, LeRoith D. Obesity and Diabetes: The Increased Risk of Cancer and Cancer-Related Mortality. Physiol Rev. 2015;95(3):727–48.

10. Godsland IF. Insulin resistance and hyperinsulinaemia in the development and progression of cancer. Clin Sci (Lond). 2009;118(5):315–32.

11. Arcidiacono B, Iiritano S, Nocera A, Possidente K, Nevolo MT, Ventura V, et al. Insulin resistance and cancer risk: an overview of the pathogenetic mechanisms. Exp Diabetes Res. 2012;2012:789174.

12. Anni M.Y. Zhang EAW, Janel L. Kopp, James D. Johnson. Hyperinsulinemia in obesity, inflammation, and cancer. Diabetes & Metabolism Journal. In press.

13. Tsujimoto T, Kajio H, Sugiyama T. Association between hyperinsulinemia and increased risk of cancer death in nonobese and obese people: A population-based observational study. Int J Cancer. 2017;141(1):102–11.

14. Perseghin G, Calori G, Lattuada G, Ragogna F, Dugnani E, Garancini MP, et al. Insulin resistance/hyperinsulinemia and cancer mortality: the Cremona study at the 15th year of follow-up. Acta Diabetol. 2012;49(6):421–8.

15. Pisani P. Hyper-insulinaemia and cancer, meta-analyses of epidemiological studies. Arch Physiol Biochem. 2008;114(1):63–70.

16. Dugnani E, Balzano G, Pasquale V, Scavini M, Aleotti F, Liberati D, et al. Insulin resistance is associated with the aggressiveness of pancreatic ductal carcinoma. Acta Diabetol. 2016;53(6):945–56.

17. Michaud DS, Wolpin B, Giovannucci E, Liu S, Cochrane B, Manson JE, et al. Prediagnostic plasma C-peptide and pancreatic cancer risk in men and women. Cancer Epidemiol Biomarkers Prev. 2007;16(10):2101–9.

18. Stolzenberg-Solomon RZ, Graubard BI, Chari S, Limburg P, Taylor PR, Virtamo J, et al. Insulin, glucose, insulin resistance, and pancreatic cancer in male smokers. JAMA. 2005;294(22):2872–8.

19. Zhang AMY, Magrill J, de Winter TJJ, Hu X, Skovso S, Schaeffer DF, et al. Endogenous Hyperinsulinemia Contributes to Pancreatic Cancer Development. Cell Metab. 2019;30(3):403–4.

20. Soares MB, Schon E, Henderson A, Karathanasis SK, Cate R, Zeitlin S, et al. RNA-mediated gene duplication: the rat preproinsulin I gene is a functional retroposon. Mol Cell Biol. 1985;5(8):2090–103.

21. Hay CW, Docherty K. Comparative analysis of insulin gene promoters: implications for diabetes research. Diabetes. 2006;55(12):3201–13.

22. Mehran AE, Templeman NM, Brigidi GS, Lim GE, Chu KY, Hu X, et al. Hyperinsulinemia drives diet-induced obesity independently of brain insulin production. Cell Metab. 2012;16(6):723–37.

23. Deltour L, Leduque P, Blume N, Madsen O, Dubois P, Jami J, et al. Differential expression of the two nonallelic proinsulin genes in the developing mouse embryo. Proc Natl Acad Sci U S A. 1993;90(2):527–31.

24. Templeman NM, Mehran AE, Johnson JD. Hyper-Variability in Circulating Insulin, High Fat Feeding Outcomes, and Effects of Reducing Ins2 Dosage in Male Ins1-Null Mice in a Specific Pathogen-Free Facility. PLoS One. 2016;11(4):e0153280.

25. Templeman NM, Clee SM, Johnson JD. Suppression of hyperinsulinaemia in growing female mice provides long-term protection against obesity. Diabetologia. 2015;58(10):2392–402.

26. Tuveson DA, Shaw AT, Willis NA, Silver DP, Jackson EL, Chang S, et al. Endogenous oncogenic K-ras(G12D) stimulates proliferation and widespread neoplastic and developmental defects. Cancer Cell. 2004;5(4):375–87.

27. Pan FC, Bankaitis ED, Boyer D, Xu X, Van de Casteele M, Magnuson MA, et al. Spatiotemporal patterns of multipotentiality in Ptf1a-expressing cells during pancreas organogenesis and injury-induced facultative restoration. Development. 2013;140(4):751–64.

28. Lee AYL, Dubois CL, Sarai K, Zarei S, Schaeffer DF, Sander M, et al. Cell of origin affects tumour development and phenotype in pancreatic ductal adenocarcinoma. Gut. 2019;68(3):487–98.

29. Matthew D Young SB. SoupX removes ambient RNA contamination from droplet-based single-cell RNA sequencing data. GigaScience. 2020;9(12).

30. Butler A, Hoffman P, Smibert P, Papalexi E, Satija R. Integrating single-cell transcriptomic data across different conditions, technologies, and species. Nat Biotechnol. 2018;36(5):411–20.

31. Stuart T, Butler A, Hoffman P, Hafemeister C, Papalexi E, Mauck WM, 3rd, et al. Comprehensive Integration of Single-Cell Data. Cell. 2019;177(7):1888–902 e21.

32. Kuleshov MV, Jones MR, Rouillard AD, Fernandez NF, Duan Q, Wang Z, et al. Enrichr: a comprehensive gene set enrichment analysis web server 2016 update. Nucleic Acids Res. 2016;44(W1):W90–7.

33. Raudvere U, Kolberg L, Kuzmin I, Arak T, Adler P, Peterson H, et al. g:Profiler: a web server for functional enrichment analysis and conversions of gene lists (2019 update). Nucleic Acids Res. 2019;47(W1):W191–W8.

34. Jassal B, Matthews L, Viteri G, Gong C, Lorente P, Fabregat A, et al. The reactome pathway knowledgebase. Nucleic Acids Res. 2020;48(D1):D498–D503.

35. Reimand J, Isserlin R, Voisin V, Kucera M, Tannus-Lopes C, Rostamianfar A, et al. Pathway enrichment analysis and visualization of omics data using g:Profiler, GSEA, Cytoscape and EnrichmentMap. Nat Protoc. 2019;14(2):482–517.

36. Kopp JL, von Figura G, Mayes E, Liu FF, Dubois CL, Morris JPt, et al. Identification of Sox9-dependent acinar-to-ductal reprogramming as the principal mechanism for initiation of pancreatic ductal adenocarcinoma. Cancer Cell. 2012;22(6):737–50.

37. Templeman NM, Skovso S, Page MM, Lim GE, Johnson JD. A causal role for hyperinsulinemia in obesity. J Endocrinol. 2017;232(3):R173–R83.

38. Templeman NM, Flibotte S, Chik JHL, Sinha S, Lim GE, Foster LJ, et al. Reduced Circulating Insulin Enhances Insulin Sensitivity in Old Mice and Extends Lifespan. Cell Rep. 2017;20(2):451–63.

39. Seifert GJ, Sander KC, Richter S, Wittel UA. Murine genotype impacts pancreatitis severity and systemic inflammation: An experimental study. Ann Med Surg (Lond). 2017;24:8–14.

40. Basturk O, Hong SM, Wood LD, Adsay NV, Albores-Saavedra J, Biankin AV, et al. A Revised Classification System and Recommendations From the Baltimore Consensus Meeting for Neoplastic Precursor Lesions in the Pancreas. Am J Surg Pathol. 2015;39(12):1730–41.

41. Elyada E, Bolisetty M, Laise P, Flynn WF, Courtois ET, Burkhart RA, et al. Cross-Species Single-Cell Analysis of Pancreatic Ductal Adenocarcinoma Reveals Antigen-Presenting Cancer-Associated Fibroblasts. Cancer Discov. 2019;9(8):1102–23.

42. Hosein AN, Huang H, Wang Z, Parmar K, Du W, Huang J, et al. Cellular heterogeneity during mouse pancreatic ductal adenocarcinoma progression at single-cell resolution. JCI Insight. 2019;5.

43. Schlesinger Y, Yosefov-Levi O, Kolodkin-Gal D, Granit RZ, Peters L, Kalifa R, et al. Single-cell transcriptomes of pancreatic preinvasive lesions and cancer reveal acinar metaplastic cells’ heterogeneity. Nat Commun. 2020;11(1):4516.

44. Chen Z, Downing S, Tzanakakis ES. Four Decades After the Discovery of Regenerating Islet-Derived (Reg) Proteins: Current Understanding and Challenges. Front Cell Dev Biol. 2019;7:235.

45. Mukherjee S, Vaishnava S, Hooper LV. Multi-layered regulation of intestinal antimicrobial defense. Cell Mol Life Sci. 2008;65(19):3019–27.

46. Zhang MY, Wang J, Guo J. Role of Regenerating Islet-Derived Protein 3A in Gastrointestinal Cancer. Front Oncol. 2019;9:1449.

47. Yin G, Du J, Cao H, Liu X, Xu Q, Xiang M. Reg3g Promotes Pancreatic Carcinogenesis in a Murine Model of Chronic Pancreatitis. Dig Dis Sci. 2015;60(12):3656–68.

48. Li Q, Wang H, Zogopoulos G, Shao Q, Dong K, Lv F, et al. Reg proteins promote acinar-to-ductal metaplasia and act as novel diagnostic and prognostic markers in pancreatic ductal adenocarcinoma. Oncotarget. 2016;7(47):77838–53.

49. Ramkissoon R, Gardner TB. Pancreatic steatosis: an update. Curr Opin Gastroenterol. 2019;35(5):440–7.

50. Gukovskaya AS, Gukovsky I. Which way to die: the regulation of acinar cell death in pancreatitis by mitochondria, calcium, and reactive oxygen species. Gastroenterology. 2011;140(7):1876–80.

51. Teper Y, Eibl G. Pancreatic Macrophages: Critical Players in Obesity-Promoted Pancreatic Cancer. Cancers (Basel). 2020;12(7).

52. Chang JH, Jiang Y, Pillarisetty VG. Role of immune cells in pancreatic cancer from bench to clinical application: An updated review. Medicine (Baltimore). 2016;95(49):e5541.

53. Clark CE, Hingorani SR, Mick R, Combs C, Tuveson DA, Vonderheide RH. Dynamics of the immune reaction to pancreatic cancer from inception to invasion. Cancer Res. 2007;67(19):9518–27.

54. Wachsmann MB, Pop LM, Vitetta ES. Pancreatic ductal adenocarcinoma: a review of immunologic aspects. J Investig Med. 2012;60(4):643–63.

55. Zhang Y, Yan W, Mathew E, Bednar F, Wan S, Collins MA, et al. CD4+ T lymphocyte ablation prevents pancreatic carcinogenesis in mice. Cancer Immunol Res. 2014;2(5):423–35.

56. Tsai S, Clemente-Casares X, Zhou AC, Lei H, Ahn JJ, Chan YT, et al. Insulin Receptor-Mediated Stimulation Boosts T Cell Immunity during Inflammation and Infection. Cell Metab. 2018;28(6):922–34 e4.

57. Covarrubias AJ, Aksoylar HI, Horng T. Control of macrophage metabolism and activation by mTOR and Akt signaling. Semin Immunol. 2015;27(4):286–96.

58. Huber M, Brehm CU, Gress TM, Buchholz M, Alashkar Alhamwe B, von Strandmann EP, et al. The Immune Microenvironment in Pancreatic Cancer. Int J Mol Sci. 2020;21(19).

59. Zhang Y, Lazarus J, Steele NG, Yan W, Lee HJ, Nwosu ZC, et al. Regulatory T-cell Depletion Alters the Tumor Microenvironment and Accelerates Pancreatic Carcinogenesis. Cancer Discov. 2020;10(3):422–39.

60. Zhang Y, Velez-Delgado A, Mathew E, Li D, Mendez FM, Flannagan K, et al. Myeloid cells are required for PD-1/PD-L1 checkpoint activation and the establishment of an immunosuppressive environment in pancreatic cancer. Gut. 2017;66(1):124–36.

61. Liu Q, Li Y, Niu Z, Zong Y, Wang M, Yao L, et al. Atorvastatin (Lipitor) attenuates the effects of aspirin on pancreatic cancerogenesis and the chemotherapeutic efficacy of gemcitabine on pancreatic cancer by promoting M2 polarized tumor associated macrophages. J Exp Clin Cancer Res. 2016;35:33.

62. Liou GY, Doppler H, Necela B, Edenfield B, Zhang L, Dawson DW, et al. Mutant KRAS-induced expression of ICAM-1 in pancreatic acinar cells causes attraction of macrophages to expedite the formation of precancerous lesions. Cancer Discov. 2015;5(1):52–63.

63. Yang S, Liu Q, Liao Q. Tumor-Associated Macrophages in Pancreatic Ductal Adenocarcinoma: Origin, Polarization, Function, and Reprogramming. Front Cell Dev Biol. 2020;8:607209.

64. Liou GY, Bastea L, Fleming A, Doppler H, Edenfield BH, Dawson DW, et al. The Presence of Interleukin-13 at Pancreatic ADM/PanIN Lesions Alters Macrophage Populations and Mediates Pancreatic Tumorigenesis. Cell Rep. 2017;19(7):1322–33.

65. Kaur K, Chang HH, Topchyan P, Cook JM, Barkhordarian A, Eibl G, et al. Deficiencies in Natural Killer Cell Numbers, Expansion, and Function at the Pre-Neoplastic Stage of Pancreatic Cancer by KRAS Mutation in the Pancreas of Obese Mice. Front Immunol. 2018;9:1229.

66. Janakiram NB, Mohammed A, Bryant T, Ritchie R, Stratton N, Jackson L, et al. Loss of natural killer T cells promotes pancreatic cancer in LSL-Kras(G12D/+) mice. Immunology. 2017;152(1):36–51.

67. Pylayeva-Gupta Y, Das S, Handler JS, Hajdu CH, Coffre M, Koralov SB, et al. IL35-Producing B Cells Promote the Development of Pancreatic Neoplasia. Cancer Discov. 2016;6(3):247–55.

68. Gunderson AJ, Kaneda MM, Tsujikawa T, Nguyen AV, Affara NI, Ruffell B, et al. Bruton Tyrosine Kinase-Dependent Immune Cell Cross-talk Drives Pancreas Cancer. Cancer Discov. 2016;6(3):270–85.

69. Spear S, Candido JB, McDermott JR, Ghirelli C, Maniati E, Beers SA, et al. Discrepancies in the Tumor Microenvironment of Spontaneous and Orthotopic Murine Models of Pancreatic Cancer Uncover a New Immunostimulatory Phenotype for B Cells. Front Immunol. 2019;10:542.

70. Strobel O, Dor Y, Alsina J, Stirman A, Lauwers G, Trainor A, et al. In vivo lineage tracing defines the role of acinar-to-ductal transdifferentiation in inflammatory ductal metaplasia. Gastroenterology. 2007;133(6):1999–2009.

