## Supplementary figures and images for "Effects of hyperinsulinemia on pancreatic cancer development and the immune microenvironment revealed through single-cell transcriptomics"

### Supplemental Figure 1

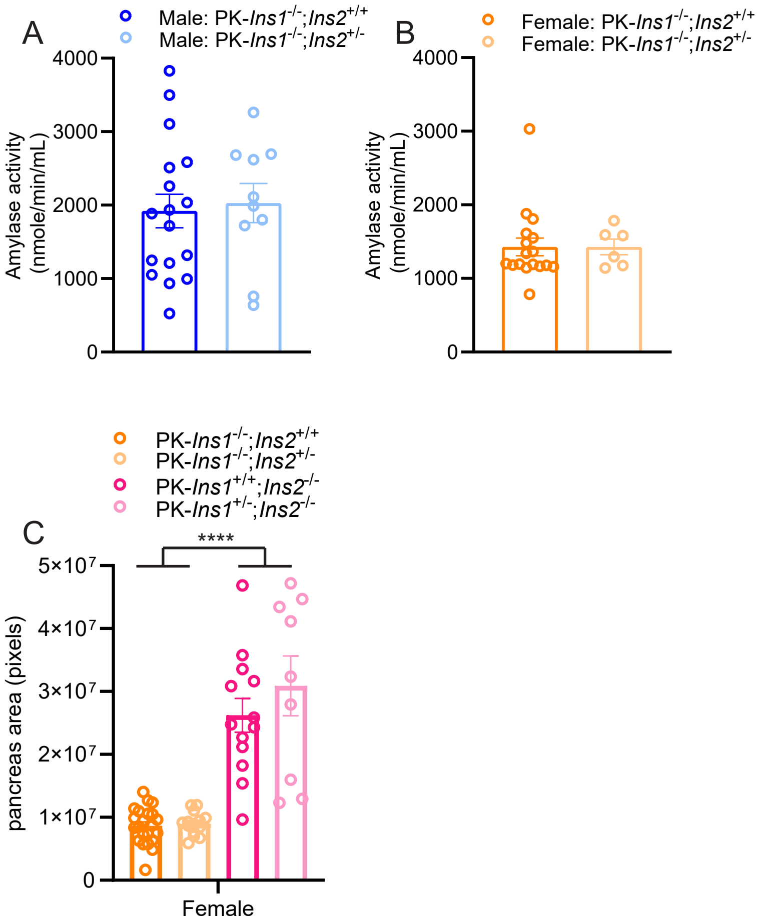


Supplemental Figure 1

### Supplemental Figure 2

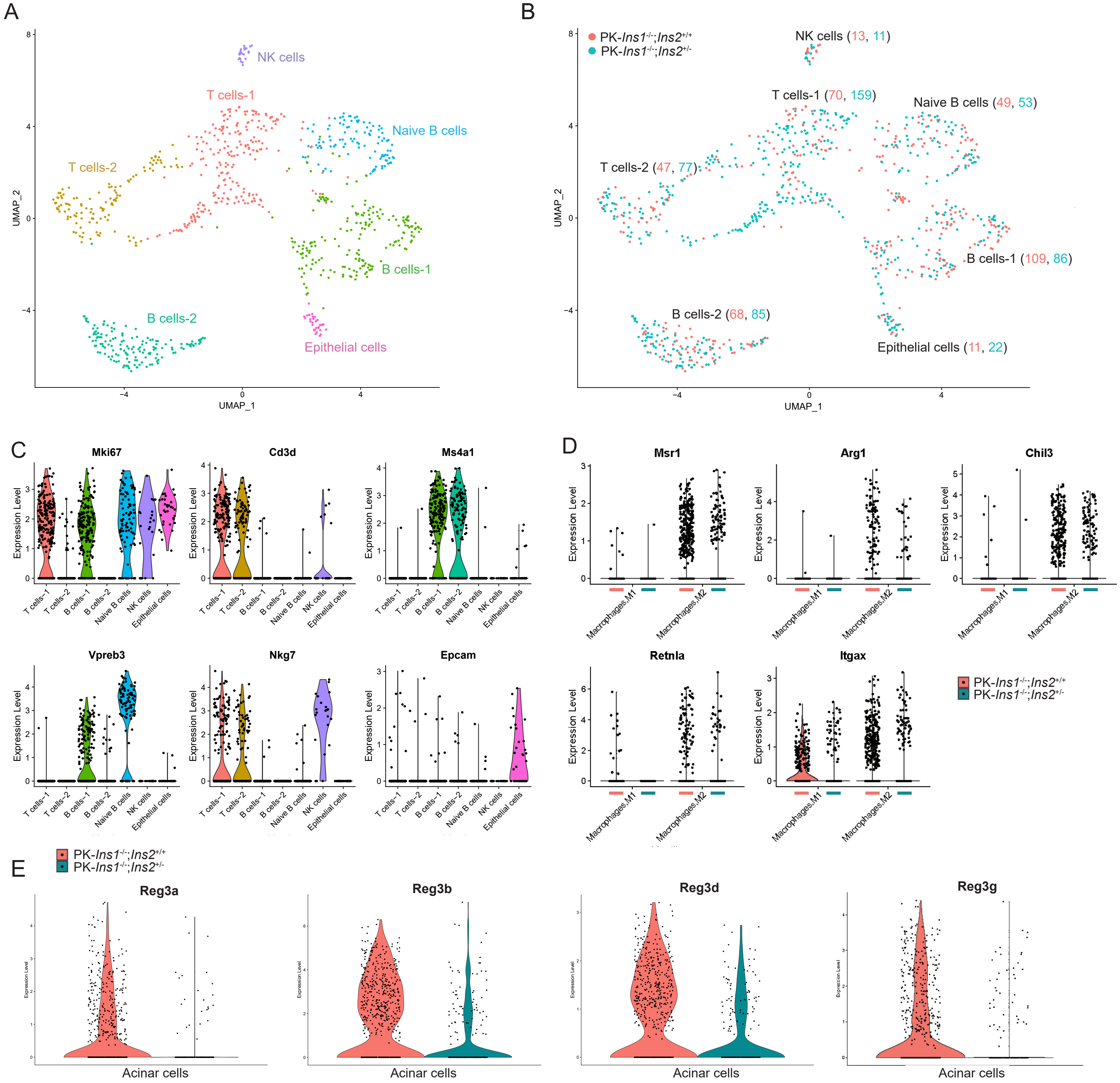
